# Geometric Analysis of Regime Shifts in Coral Reef Communities

**DOI:** 10.1101/2020.01.10.899179

**Authors:** Edward W. Tekwa, Lisa C. McManus, Ariel Greiner, Madhavi A. Colton, Michael S. Webster, Malin L. Pinsky

**Author notes:** Correspondence to: Edward W. Tekwa.

## Abstract

Coral reefs are among the many communities believed to exhibit regime shifts between alternative stable states, single-species dominance, and coexistence. Proposed drivers of regime shifts include changes in grazing, spatial clustering, and ocean temperature. Here we distill the dynamic regimes of coral-macroalgal interaction into a three-dimensional geometry, akin to thermodynamic phase diagrams of state transitions, to facilitate analysis. Specific regime-shifting forces can be understood as bifurcation vectors through the cubic regime geometry. This geometric perspective allows us to understand multiple forces simultaneously in terms of the stability and persistence of interacting species. For example, in a coral-macroalgae community, grazing on macroalgae can lead to alternative stable states when there is no spatial clustering (e.g., high habitat connectivity). However, with spatial clustering, grazing can lead to coexistence because of elevated local intraspecific competition. The geometrical analysis of regime shifts is applicable to any two-species communities and can help conservation efforts navigate complexity and abrupt changes.

## I. Introduction

Regime shifts and alternative stable states have been implicated in many communities, including coral reefs (Hughes et al. 2017), shallow lakes (Scheffer et al. 1993), kelp beds (Ling et al. 2014), and terrestrial forests (Hirota et al. 2011). Discontinuous shifts in community dynamics due to gradual environmental changes imply that conservation and management may have to anticipate and confront historical legacy traps (Scheffer et al. 2001, Tekwa et al. 2019a). The potential for regime shifts is a pressing concern in the Anthropocene, as exemplified by recent heat waves driving coral reefs to a depauperate state (Hughes et al. 2019). Coral reefs have been intensely studied and share general features with a wide range of other communities suggested to exhibit regime shifts, particularly those that feature two species whose interactions are selectively mediated by grazers, nutrients, fire, or temperature (Mumby et al. 2007, Staver and Levin 2012, Graham et al. 2015, Schmitt et al. 2019). However, there remains disagreement about the evidence for regime shifts and alternative stable states among coral reefs (Bruno et al. 2009, Dudgeon et al. 2010, Mumby et al. 2013) and other communities (Schröder et al. 2005). One possible explanation for this disagreement is that there are different mechanisms leading to regime shifts even within one ecosystem type such as coral reefs (van de Leemput et al. 2016), such that empirical examinations focusing on one mechanism will yield negative results across sites.

In the coral reef literature, multiple regime shift mechanisms have been modelled separately, including interspecific competition among coral species, interspecific competition between coral and macroalgae, predator-prey interaction, and grazer-mediated interaction (Knowlton 1992, Mumby et al. 2007, Petraitis and Hoffman 2010, van de Leemput et al. 2016). These mechanisms hinge on space being a limiting resource for benthic coral reef communities (McCook et al. 2001, Sandin and McNamara 2012), as is evident by the common use of coral cover (maximum of 100%) in the literature (Jokiel et al. 2015). However, models that track coral cover often treat space as if it were any other limiting non-spatial resource, without explicitly incorporating spatial dynamics (Elmhirst et al. 2009, Anthony et al. 2011, Blackwood et al. 2011, Baskett et al. 2014, Fabina et al. 2015, McManus et al. 2019). However, we know from the broader ecological literature that spatial clustering, arising from low habitat connectivity or limited dispersal, can strongly determine species stability in communities even with linear interaction responses (Bolker and Pacala 1999, Chesson 2000). There is therefore a need to synthesize the variety of spatial and non-spatial mechanisms of coral reef regime shifts in general ecological terms.

Here we propose simple modifications to a bi-linear mathematical model (Volterra 1926, Lotka 1978, Neuhauser and Pacala 1999) so as to use generic community ecological terms to synthesize spatial, temperature, and grazing effects on coral macroalgal interactions. This model reveals the basic ingredients that lead to alternative stable states or coexistence of corals and macroalgae on coral reefs, as well as what these species stability outcomes mean for the aggregate community. We then distill the model to three parameters that completely define the possible dynamic regimes and that can be visualized as a cubic volume. We show how previously suggested bifurcating factors—such as grazing, spatial clustering, and warming—are different vectors traversing this cubic parameter space. The ultimate goal of this formalism is the identification of generic bifurcation dimensions (local competition and intrinsic growth metrics) that will allow scientists and conservation managers to generate and test hypotheses regarding the presence or absence of regime shifts without narrowly focusing on single region-or system-specific mechanisms.

## II. Methods

We first present the Lotka-Volterra model as a foundation for two-species interactions, then show that a coral-macroalgae model can be analyzed as a special case and extended to incorporate temperature dependence. We then incorporate spatial clustering into the models, arriving at a general Spatial Lotka-Volterra formulation of dynamic regimes in two-species systems. Finally, we add temperature dependent growth. The specific spatial and temperature-dependence introduced for coral-macroalgal interactions allow us to subsequently explore how grazing, spatial clustering, and warming affect coral reef communities’ dynamic regimes.

### Lotka-Volterra Model

We first restate the classic two-species competitive Lotka-Volterra equations and their well-known implications for bistability and coexistence (Volterra 1926, Lotka 1978). The species in these equations can represent coral and macroalgae. The Lotka-Volterra model assumes that each species has intrinsic growth rate (*r_i_*) and mortality (*m_i_*). In addition, competition between species *i* and *j* results in linear per-capita growth rate changes (-*r_i_a_ij_*) that scale with the density of the other species (*N_j_*) (Table 1). There are three non-trivial equilibria sets, including species 1 dominance (case 1), species 2 dominance (case 2), and coexistence (Table A1). Stability analysis (Appendix: Lotka-Volterra Model) shows that the single-species equilibrium for species *i* is stable if:

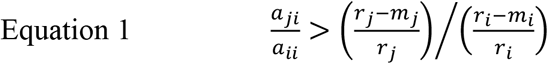

**Table 1.** Model equations. The dynamic equations are given in the form of *dN_i_*/(*N_i_dt*)=Σ(*coefficient* × *state*) where each coefficient is highlighted in orange and the corresponding state is the bracketed variable given in the header row. Subscript *i* refers to the focal species and *j≠i*. All symbols are defined in Table 2.

| Model | Species (i) | Density changes | Intrinsic rate ($\times 1$) | Intraspecific interaction ($\times N_i$) | Interspecific interaction ($\times N_j$) | Higher-order interspecific interaction ($\times (N_j^2 + N_j^3 + \dots)$) |
| --- | --- | --- | --- | --- | --- | --- |
| Lotka-Volterra | 1 | $\frac{dN_1}{N_1 dt} = \Sigma$ | $r_1 - m_1$ | $-r_1 a_{11}$ | $-r_1 a_{12}$ | 0 |
| | 2 | $\frac{dN_2}{N_2 dt} = \Sigma$ | $r_2 - m_2$ | $-r_2 a_{22}$ | $-r_2 a_{21}$ | 0 |
| Mumby model | 1 coral | $\frac{dN_1}{N_1 dt} = \Sigma$ | $r - d$ | $-r$ | $-(r + a)$ | 0 |
| | 2 algae | $\frac{dN_2}{N_2 dt} = \Sigma$ | $\gamma - g$ | $-\gamma$ | $-(\gamma + g - a)$ | $-g$ |
| Spatial Lotka-Volterra | 1 | $\frac{dN_1}{N_1 dt} = \Sigma$ | $r_1 - m_1$ | $-r_1 a_{11} C_{11}$ | $-r_1 a_{12} C_{12}$ | 0 |
| | 2 | $\frac{dN_2}{N_2 dt} = \Sigma$ | $r_2 - m_2$ | $-r_2 a_{22} C_{22}$ | $-r_2 a_{21} C_{12}$ | 0 |

That is, if the ratio of interspecific competition (of species *j* on *i, a_ji_*) over intraspecific competition (of *i, a_ii_*) is greater than the ratio of species *j*’s isolated equilibrium density ((*r_j_-m_j_*)/*r_j_*) over species *i*’s isolated equilibrium density ((*r_i_-m_i_*)/*r_i_*) (when intraspecific competitions are equal, *a*_11_=*a*_22_), then the dominance of species *i* (with *j* locally extirpated) is stable. If the condition in Equation 1 is true for only *i*=1 but not *i*=2, then species 1 competitively excludes species 2 deterministically, and vice versa for species 2 competitively excluding species 1. If the condition is false for both species, then coexistence is stable. However, if Equation 1 is true for *i*=1 and for *i*=2, then coexistence is unstable and alternative stable states occur, with either species dominating depending on initial conditions.

### Coral-Macroalgae Model

We next we show that models based on the Lotka-Volterra formulation can help understand competitive exclusion, bistability, and coexistence conditions in prominent coral-macroalgae models. The Mumby model (Mumby et al. 2007) and related models (Li et al. 2014) consider coral (*N*_1_) and macroalgal (*N*_2_) cover. These models exhibit bistability when an implicit herbivore’s grazing rate on macroalgae (*g*) is at an intermediate value. The Mumby model can be rewritten in Lotka-Volterra form, with terms arranged according to intraspecific and interspecific interactions (Table 1, Appendix: Coral-Macroalgae Model).

With this formulation, it becomes clear that the Mumby model is a particular specification of the Lotka-Volterra model in which grazing reduces the intrinsic growth rate of and increases the interspecific competition on macroalgae. This formulation also reveals the implicit assumptions about competition, namely that interspecific competition is greater than intraspecific competition for corals under any grazing rate. Interspecific competition is also greater than intraspecific competition for macroalgae when grazing rate is sufficiently high (Appendix, Table 1). Thus, the alternative stable states observed in the model can be understood in terms of the Lotka-Volterra terminology of interspecific versus intraspecific competition (Equation 1).

In addition, the Mumby model features a negative grazing effect on macroalgae that increases in magnitude geometrically with coral cover (*N*_1_^2^+ *N*_1_^3^+…+*N*_1_^∞^) (Appendix, Table 1). Dropping these higher-order interactions shrinks but does not eliminate the bistable region and, in fact, the alternative stable states remain identical (Figure A1, Equation 10, see Table 2 for parameter values). Therefore, the Lotka-Volterra model appears sufficiently nuanced to represent alternative stable state dynamics between coral and macroalgae.

**Table 2.** Symbol definitions. Parameter values are for Figures 2–4.

| Definition | Species 1 (coral)<br>parameter values | Species 2 (macroalgae)<br>parameter values |
| --- | --- | --- |
| ***Coral-Macroalgae Model*** (without spatial + temperature dependence) |  |  |
| macroalgal overgrowth on coral | $a=1.1$ | |
| coral mortality | $d=0.5$ | |
| grazing rate | | $g=[0.55 - 0.85]$ |
| birth rate | $r=1$ | $\gamma=1.1$ |
| ***Lotka-Volterra Equivalent*** (with spatial + temperature dependence) |  |  |
| intraspecific interaction effect | $a_{11}=C_{ii}$ | $a_{22}=C_{ii}$ |
| interspecific interaction effect | $a_{12}=(r_1+a)C_{12}/r_1$ | $a_{21}=(r_2+g-a)C_{12}/r_2$ |
| relative (intra-to-inter) clustering | $C_{ii}/C_{ij}=[1, 2, 4]$ | |
| intraspecific clustering | $C_{ii}=[1, 1.19, 1.41]$ | |
| mortality | $m_1=d$ | $m_2=g$ |
| density or cover | $0 \leq N_i \leq 1$ | |
| intrinsic growth rate | $r \cdot \exp(-\Delta T^2/(2\sigma_1^2))$ | $\gamma \cdot \exp(-\Delta T^2/(2\sigma_2^2))$ |
| thermal tolerance | $\sigma_1=1$ | $\sigma_1=\sqrt{2}$ |
| actual - optimal temperature | $\Delta T=[0, 1]$ | |

We note that Lotka-Volterra-based models traditionally define species state (*N_i_*) as density (biomass or abundance per area), while the coral literature tracks proportion of habitat covered by biomass (maximum of one or 100%) (Jokiel et al. 2015). Given any arbitrary area unit, density in the Lotka-Volterra model can also be set to a maximum of one both locally and globally by adjusting the competition coefficients *a_ij_*. Thus, density and percent cover are interchangeable for the subsequent results.

Having established the connection between the Lotka-Volterra model and the Mumby model, we now proceed to incorporate space into the Lotka-Volterra model.

### Spatial Lotka-Volterra Model

Spatial competition is an implicit assumption in the coral-macroalgal interaction (McCook et al. 2001, Sandin and McNamara 2012). Here we explicitly consider how spatial dynamics affect coral and macroalgae using the Lotka-Volterra formulation. The Lotka-Volterra model can be changed into a spatial version using the spatial moment framework (Durrett and Levin 1994, Bolker and Pacala 1999, Lion and Baalen 2008, Tekwa et al. 2015). According to the spatial moment framework, interaction neighbour densities for a focal species *i* in a non-spatial model (*N_j_*) can be replaced by the local density *N_ij_*, or *C_ij_N_j_* (related to the second spatial moment, see Appendix) where *C_ij_* is a continuous-space clustering coefficient. This clustering coefficient is relevant across a variety of ways of thinking about space, including continuous space (with neighbours weighted by distance), discrete space such as habitat networks or metacommunities (with neighbours being within a patch), or social networks (with neighbours being connected nodes) (Lion and Baalen 2008, Tekwa et al. 2017). *N_ij_* or *C_ij_N_j_* expresses the average number of species *j* neighbours that an individual of species *i* interacts with per area per time, and can be different from *N_j_*, the average number of neighbours that an individual would interact with if all were randomly distributed or if the interaction neighbourhood were the entire community (Figure 1). In network terminology with two species, *N_ii_* is the average node degree in the within-species network, whereas *N_ij_* (*i≠j*) is the average node degree in the bipartite network (where the links are between species).

**Figure 1.**
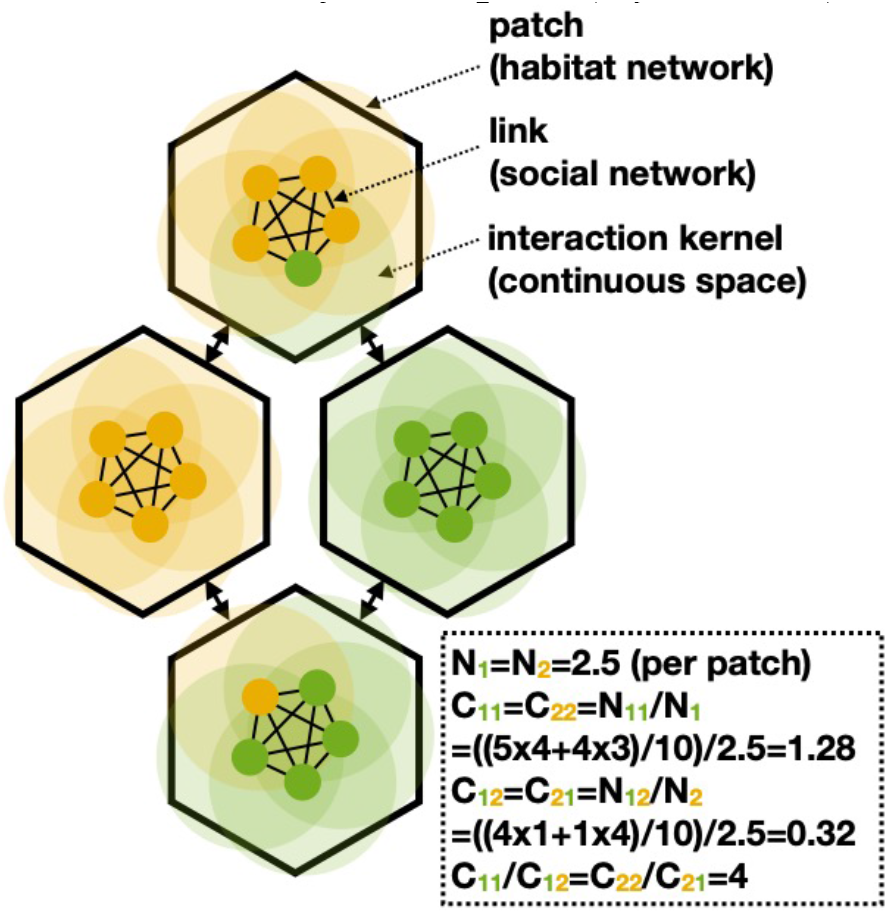
Descriptions of spatial clustering. The spatial clustering of individuals (circles) of two species can be conceptualized in three different ways. First, patches (hexagons) in a habitat network can delimit which individuals are interaction neighbours. Second, links (thin lines) in a social network can specify which pair of individuals interact at a given time. Third, interaction kernels (circular shades) can weigh individuals within a certain distance as neighbours. The spatial clustering discussed in the main text can be described under any of these three frameworks with continuous-space clustering coefficients *C_ij_*. These coefficients can be tallied in terms of the average number of neighbours (or node degree) *j* that *i* experiences (*N_ij_*) and the global average number of individuals *i* per area or patch (*N_i_*). Sample calculations of *N_i_*, *N_ij_*, *C_ij_*, and relative clustering (*C_ii_*/*C_ij_*) are obtained by taking averages and ratios of individual and neighbour counts (see box).

The clustering coefficient is convenient because it captures spatial clustering effects as a single multiplicative factor, indicating how many more (when *C_ij_*> 1) or fewer (when *C_ij_*<1) times an individual of species *i* encounters an individual of species *j* than the global density of *j*. The higher the value of *C_ij_*, the more clustered *j* is around *i*. This also allows one to write an interaction effect on population growth rate (*dN_i_/N_i_dt*) as *a_ij_C_ij_N_j_*. In this form, it is clear that the dynamic equations are the same as the non-spatial Lotka-Volterra equations, with interaction coefficients *a_ij_* replaced by *a_ij_C_ij_*. That is, spatial clustering scales up the effective interaction effects. By definition, *C_ij_*=*C_ji_* (Tekwa et al. 2015). Spatial clustering can be due to either endogenous (low dispersal and pattern formation) or exogenous (habitat connectivity and matrix constraint) processes. In particular, low dispersal leads to *C_ij_* being greater than one within species (*C_ii_*>1) and less than one between species (*C_ij_*<1) because offspring tend to be near parents (Bolker and Pacala 1999, Lion and Baalen 2008, Tekwa et al. 2019b). Here we assume that clustering is constant through time and ignore possible dependency on *N_i_* or higher moments (Bolker and Pacala 1999). Among species or morphs that are very similar, as in an incremental evolutionary process without population size dynamics, it has been shown that relative clustering (*C_i_i*/*C_ij_*) is constant (Tarnita et al. 2009, Nathanson et al. 2009, Tekwa et al. 2015). In the more general ecological case where species can be very different, more habitat connectivity or higher movement rates are still expected to create less relative clustering (approaching one with the highest connectivity or movement rates) (Bolker and Pacala 1997, Tarnita et al. 2009, Tekwa et al. 2019b). Thus, the constant clustering assumption is an approximation that should roughly capture spatial effects on regime dynamics.

Spatial clustering affects coral and macroalgal competition terms under the Spatial Lotka-Volterra framework. By matching terms in the Spatial Lotka-Volterra model and the coral-macroalgae model (Table 1), we find that intraspecific competition is 1 without spatial clustering, and increases with within-species clustering (*C_ii_*, Table 2). Interspecific competition effects, on the other hand, are moderated by both space (*C_ij_*, Table 2).

With spatial considerations the stability criterion for species *i* dominance becomes:

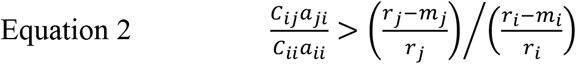

This inequality is harder to attain when relative clustering (*C_ii_*/*C_ij_*) is high. Thus, clustering can lead to global coexistence, even when locally there tends to be one or the other species dominating. The finding is congruent with the well-known hypothesis that spatial variation promotes coexistence (Chesson 2000).

### Temperature Dependence

Warming is recognized as one of the most dramatic factors affecting coral reefs (Hughes et al. 2019). As a simple and analytically tractable way to consider temperature, we assume that intrinsic growth rates *r_i_* are maximal when temperature matches the historical temperature (*r* for corals and *γ* for macroalgae), and that growth rates decrease when temperature deviates from these optima according to (non-standardized) Gaussian functions. A species’ thermal tolerance is the standard deviations of the Gaussian function. Further, we assume that macroalgae have a wider thermal tolerance (*σ*_1_) than corals (*σ*_2_, Table 2). Mortality rates are assumed constant in temperature for corals (*d*) and for macroalgae (*g*).

## III. Results

We use stability criteria in the spatial Lotka-Volterra model to show how dynamic regimes in two-species (e.g., coral-macroalgal) communities can be generically described using simple geometry with only three parameters for competition and growth. We then show how the effects of grazing, spatial clustering, and warming translate to changes in these three competition and growth parameters to affect dynamic outcomes in the coral-macroalgal system. We aim to show that diverse mechanisms of community regime shifts can be synthesized under a common, low-dimensional geometric framework.

### Geometry of Dynamic Regimes

The community dynamic regimes of a two-species spatial Lotka-Volterra model are determined by two inequalities involving three parameters. From Equation 2, the three parameters are 1) the local species 1 intra-to-interspecific cross competition ratio *α_1_*; 2) the local species 2 intra-to-interspecific cross competition ratio *α_2_*; and 3) the intrinsic growth inequality ratio between species 2 and 1, *f_21_* (see Table 3). The competition ratios are called “cross competition”, because they are ratios of the intraspecific competition effect on the focal species relative to the interspecific competition effect on the other species. Competition ratios also encapsulate the effect of spatial clustering, which is positive and multiplicative. Table 3 shows that the possible combinations of inequalities produce the four dynamic regimes of alternative stable states, species 1 only, species 2 only, and coexistence. The points where the three parameters coincide (1/*α_1_*=*α_2_*=*f_21_*), for example at *α_1_*= *α_2_*=*f_21_*=1, are “quadruple points” where the four dynamic regimes collide (named after the triple point in the thermodynamic phases of solid, liquid, and gas) (Maxwell and Harman 1990). Some illustrative bifurcation calculations are shown in Table A2 and Table A3 to demonstrate that increases in relative clustering shift dynamics from “alternative stable states” to “species 2 only” and eventually to “coexistence.” Similarly, increases in grazing shifts the dynamics from “species 2 only” to “alternative stable states” to “species 1 only.”

**Table 3.** Conditions for each community dynamic regime. The variables that determine dynamic regimes are 1) intra-to-interspecific cross competition ratio 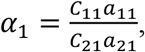, 2) intra-to-interspecific cross competition ratio 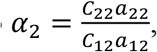, and 3) intrinsic growth inequality 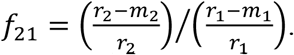.

| Conditions | Community Dynamic Regimes |
| --- | --- |
| $1/\alpha_1 > f_{21} > \alpha_2$ | alternative stable states |
| $1/\alpha_1 > f_{21} < \alpha_2$ | species 1 only |
| $1/\alpha_1 < f_{21} > \alpha_2$ | species 2 only |
| $1/\alpha_1 < f_{21} < \alpha_2$ | coexistence |

The three parameters constitute the coordinates in which the stability of each species can change. The planes 1/*α_1_* = *f_21_* and *f_21_*=*α_2_* bisect, respectively, regions where species 1 and species 2 dominance are marginally stable. In particular, in log-space these planes are flat (because all dimensions are ratios, Figure 2A). Using these planes, we construct a volume with the three dimensions as axes, and dynamic regime as categorical outcomes coded by color (Figure 2B). This cube completely describes all possible dynamic regimes and their relationships to parameters in the spatial Lotka-Volterra model.

**Figure 2.**
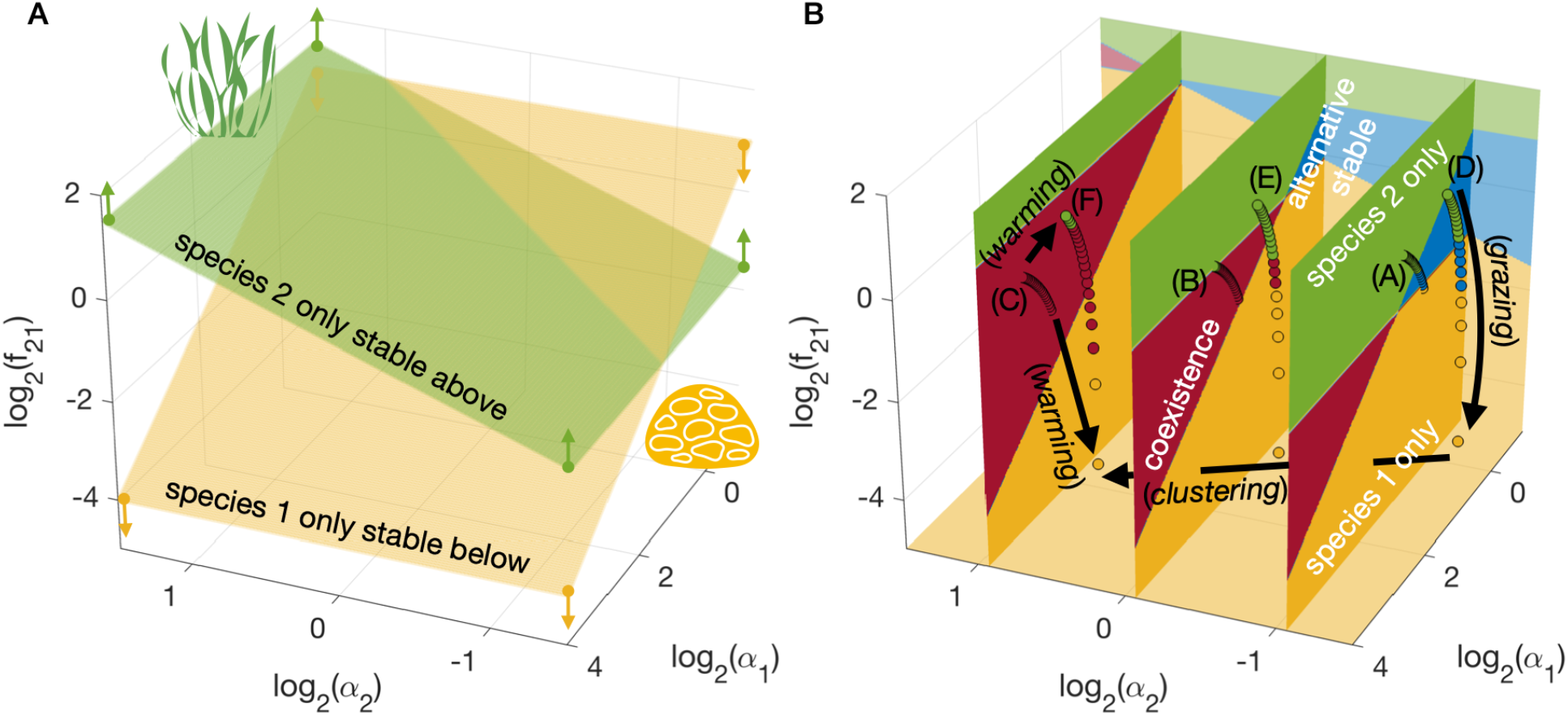
Geometric representation of the relationship between Lotka-Volterra parameters and the four possible dynamic regimes. The dimensions are the species 1 intra-to-interspecific cross-competition log-ratio (log_2_(*α_1_*)), the species 2 cross-competition log-ratio (log_2_(*α_2_*)), and the intrinsic growth log-inequality of species 2 over species 1 (log_2_(*f_21_*)). **(A)** The two-species spatial Lotka-Volterra model’s dynamic regimes are separated by two planes that define the marginal stability of each species’ dominance. These planes bisect each other and create four dynamic regimes **(B),** which are illustrated using three two-dimensional cross-sections (colored regimes with white text). Bifurcation vectors (black arrows and text) show the effects of grazing, warming, and spatial clustering. Letters A-F corresponding to subplots in Figure 3. Series of circles colored by regimes represent how equidistant increments in grazing in a coral-macroalgae model traverse the regime geometry. The series start at three different and fixed spatial clustering and two warming levels.

The dynamic regime geometry distills the spatial Lotka-Volterra model into three bifurcation dimensions that summarize competition and intrinsic growth properties (*α_1_*,*α_2_*, *f_21_*). This is a drastic dimensionality reduction from the original spatial Lotka-Volterra model (11 dimensions: *a_11_*, *a_12_*, *a_21_*, *a_22_*, *C_11_*, *C_12_*, *C_22_*, *m_1_*, *m_2_*, *r_1_*, *r_2_*) and the linearized coral-macroalgal model (5 dimensions: *a*, *d*, *g*, *γ*, *r*) (Table 1 and Table 2). The dimensionality reduction also means that there are multiple ways (multiple combinatorial changes in the original parameters) to achieve the same bifurcations. For example, equal changes to either relative clustering *C_11_/C_21_* or to the local competition ratio *a_11_/a_21_* results in the same change in *α_1_* and therefore the same sequence of regime shifts – either from coexistence to species 1 only, or from species 2 only to alternative stable states depending on *f_21_* (Figure 2B).

We focussed here on coral-macroalgal competition, but the results in this section apply to any two species by virtue of the generic spatial Lotka-Volterra formulation.

### System-Specific Outcomes

The categorization of dynamic regimes and dimensional reduction allow one to take a geometric approach to reasoning. Here we illustrate the utility and limitation of geometric reasoning by comparing it against species-level outcomes from a particular set of parameters. In this system, we explore how changes in grazing (Mumby et al. 2007), spatial clustering (Bolker and Pacala 1999), and warming (Hughes et al. 2019) affect dynamic regimes – quantities that should be obtainable from geometric reasoning alone. We also explore effects on coral and macroalgae covers or densities – quantities that are related to but are more specific than categorical regimes (see Table A4 for parameter values and numerical outcomes from this example).

First, we show how parameter changes can be represented as bifurcation vectors corresponding to the geometric coordinates of *α_1_*,*α_2_*, and *f_21_* (series of circles in Figure 2B). As grazing increases, it decreases the relative growth of macroalgae versus coral (*f_21_*) and decreases the cross-competition ratio (relative intraspecific competition) for macroalgae (*α_1_*). A major effect is to drive the system towards the lower part of Fig. 2B. In contrast, increases in spatial clustering increase the cross-competition ratios for both species (*α_1_*,*α_2_*), driving the system towards the front left corner of Fig. 2B.

The effect of warming is more complicated. Warming decreases the cross-competition ratios (*α_1_*,*α_2_*) independently from clustering and grazing. Less intuitively, warming increases the growth inequality (*f_21_*) at low grazing due to macroalgae’s wider thermal tolerance, but decreases the growth inequality at high grazing where even a slight drop in γ pushes macroalgae closer to zero growth (see Table 1). The result is an expanded range of *f_21_* values traversed by grazing variation when combined with warming.

We next compare coral and macroalgal cover changes (Figure 3) to corresponding regime shifts from the geometric perspective (Figure 2). Under no warming and no spatial clustering, increases in grazing transition the community from macroalgal dominance to alternative stable states to coral dominance (Figure 3A). With more clustering, macroalgal dominance is only realized at low grazing, and coexistence becomes more likely at high grazing (Figure 3B, C). With increased temperatures, grazing traverses a larger competition-growth parameter space and therefore its effects are magnified. The regions for macroalgae (at low grazing) or coral dominance (at high grazing) increase, and the regions for coexistence or alternative stable states decrease (Figure 3D-F) when compared to the case with baseline temperatures (Figure 3A-C). The geometrically predicted alternative stable states and coexistence regimes, corresponding to cases in Figure 3A and F, are confirmed with phase diagrams where transient trajectories with different initial conditions converge on the expected number of stable equilibria (Figure 4).

**Figure 3.**
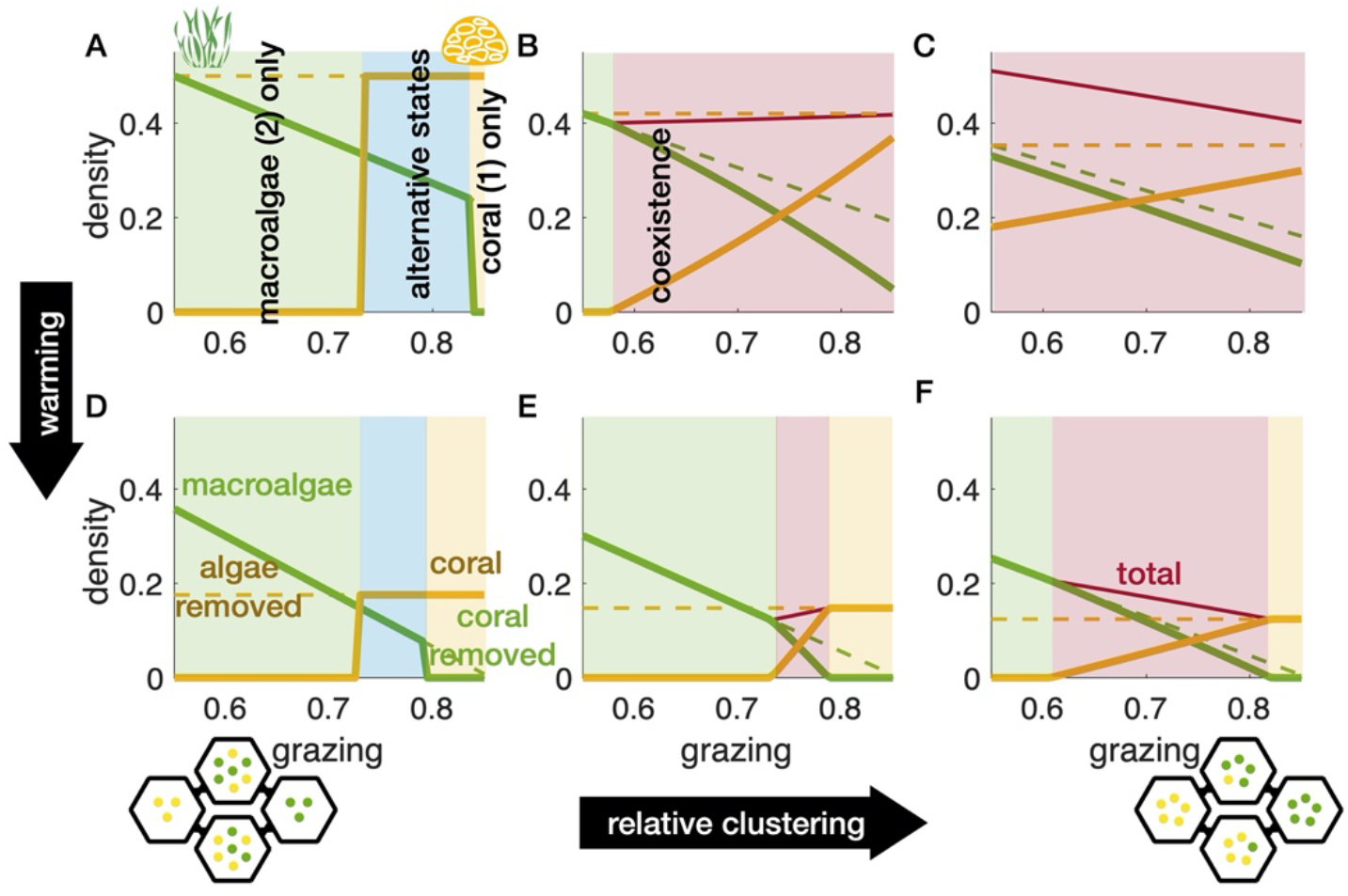
Regime shifts and coral-macroalgal density changes driven by changes in grazing. Results are from the spatial Lotka-Volterra model (see Table 2 for parameterization). Plots show macroalgal cover or density (green line), coral cover (yellow line), macroalgae or coral with the other artificially removed (dotted lines, to contrast with coexistence effects), and total cover of both taxa during coexistence (maroon line). **(A-C)** Baseline temperatures, with relative clustering (*C_ii_*/*C_ij_*) at 1, 2, or 4 (from left to right). **(D-F)** 1 ° C warming, with relative clustering being 1, 2, and 4. The shades indicate the regimes of macroalgal dominance (green), alternative stable states (blue), coral dominance (yellow), and coexistence (red). Yellow and green dots in patch diagrams at the bottom illustrate cases of low (left) versus high (right) relative clustering.

**Figure 4.**
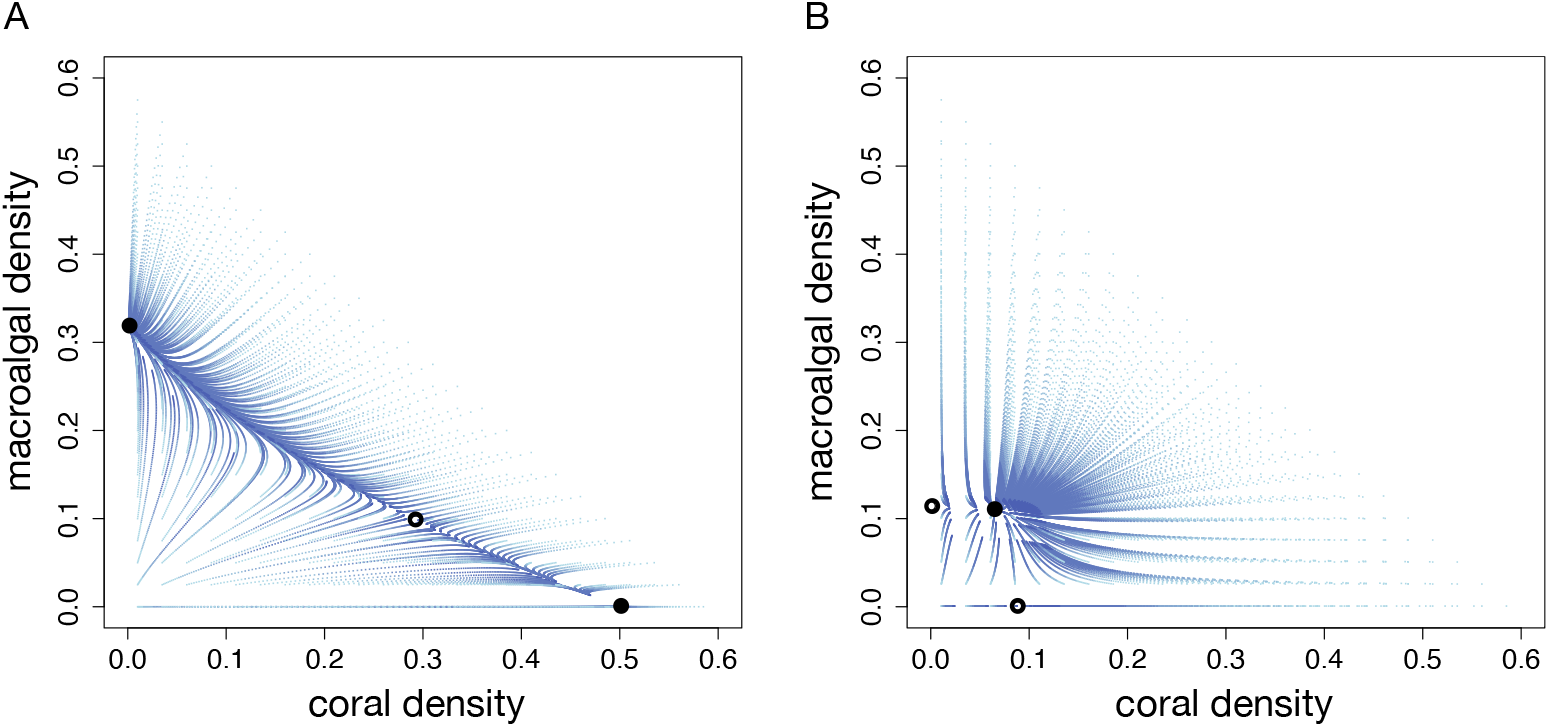
Phase diagrams of Lotka-Volterra coral-macroalgal dynamics. Trajectories (blue) are shown for 100 time steps starting at evenly spaced initial densities, with darker colour indicating densities at later times. Filled circles are analytically derived stable equilibria, while open circles are unstable equilibria. **A.** Trajectories corresponding to baseline temperatures, no spatial clustering, and a grazing rate of 0.75 showing alternative stable states (scenario in Figure 3A). **B.** Trajectories corresponding to an increased temperature, high spatial clustering, and a grazing rate of 0.75 showing coexistence (scenario in Figure 3F).

In summary, the outcomes for the specifically parameterized coral-macroalgae system illustrate levels of dynamic precision that cannot be gleaned from geometric reasoning alone; but the dynamic regime predictions from geometry remain accurate. The most detailed features of a dynamic system – transient trajectories (Figure 4) – are only partly captured by equilibrium analyses (Figure 3). Equilibria, or expected coral and macroalgal densities, are in turn not captured by regime geometry (Figure 2). Nevertheless, with only three coordinates *α_1_*,*α_2_*, and *f_21_* (Figure 2 vectors and matching color codes in Figure 3 and Table A4), regime shifts caused by multiple bifurcating forces including grazing, warming, and spatial clustering can be inferred using geometric reasoning alone (series of circles in Figure 3B).

## IV. Discussion

Regime shifts have been a focus of conservation in an era of change (Steffen et al. 2015), and coral reefs have served both as a model for understanding such shifts and as an important biome that is a focus of substantial conservation efforts (Hughes et al. 2017). Conservation efforts are, however, confounded in part by the diverse and disparate proposals for mechanisms that drive regime shifts in coral reefs (Mumby et al. 2013, van de Leemput et al. 2016, Hughes et al. 2019). Here, we provided a theoretical synthesis that captures the essential dynamics within coral reefs and other competitive communities. Further, we found that the dynamic regimes of alternative stable states, single-species dominance, and coexistence can be fully determined by only three synthetic parameters. These three parameters are a drastic dimensionality reduction, an approach that has proven useful for related studies of dynamic transitions (Jiang et al. 2018). The reduced parameter set summarizes intraspecific versus interspecific spatial competition effects (*α_1_*,*α_2_*), as well as intrinsic growth differences between species (*f_21_*). The three parameters form a cubic volume that allows for a geometric analysis of regime shifts. Ecologically realistic bifurcations or regime-shifting forces, such as grazing, spatial clustering changes, and warming, can be visualized as vectors through the dynamic regime cube.

The regime perspective produces conservation-relevant insights despite ignoring species-specific outcomes. In a coral-macroalgae system, we showed that grazing decreases the intrinsic growth difference *f_21_* and moves the system away from macroalgal dominance. Warming stretches the geometric space that grazing variation traverses, thereby increasing the likelihood of either coral or macroalgae dominating. Spatial clustering on the other hand moves the system towards higher intraspecific competition relative to interspecific competition (*α_1_* and *α_2_*), which promotes coexistence and reduces the effectiveness of grazing in inducing coral dominance. These geometric reasonings suggest that the protection of grazers will have an enhanced positive effect on coral conservation under warming in conjunction with low spatial clustering (such as, for example, by maintaining habitat connectivity between reefs). In contrast, if grazer protection fails in the face of fishing pressure (Botsford et al. 1997, Costello et al. 2016, Tekwa et al. 2019a), then high clustering through low habitat connectivity (e.g., from greater distance between protected areas) may actually enhance coral persistence through spatial coexistence mechanisms (Chesson 2000), although at much lower levels than if both grazers and habitat connectivity are protected. These geometric results illustrate that multiple management tools, such as controls on grazing and connectivity, can interact to produce conservation outcomes.

The geometry of regime shifts resembles other uses of graphical reasoning such as population growth isoclines (Tilman 1980, Knowlton 1992, McCann and Yodzis 1995) and economic phase diagrams (Gordon 1954, Solow 1956). Our approach differs due to its basis in synthetic stability criteria (see Appendix: Lotka-Volterra Model) that directly provide intuition regarding community outcomes rather than flows. The approach also focuses on how dynamic regimes shift with all possible parameter changes, in contrast to traditional Lotka-Volterra studies that often explored transient dynamics and equilibria at fixed parameterizations or variations along one parameter (Bomze 1983, Neuhauser and Pacala 1999). Regime geometry is most analogous to phase diagrams of thermodynamic states, such as the *p-v-T* (pressure-volume-temperature) diagram of a substance’s transitions between solid, liquid, and gas states (van der Waals 1873, Gibbs 1873, Verwiebe 1939, Maxwell and Harman 1990). If regime geometry and thermodynamic phase diagrams are truly analogous, then dynamics deviating from the spatial Lotka-Volterra model (nonlinear terms) could appear as modified marginal planes and regime volumes in the competition-growth space. The success of thermodynamic phase diagrams for different substances has facilitated engineering advances such as the motor and refrigeration, suggesting that regime geometry can provide a boost for conservation and ecosystem engineering by moving theoretical reasoning from mathematics to a more intuitive visualization.

The ability to geometrically represent system-specific bifurcations in generic ecological terms allows for a synthetic understanding of a wide variety of ecological communities. Regime shifts in lakes (Scheffer et al. 1993), kelp forests (Ling et al. 2014), and terrestrial forests (Hirota et al. 2011) share both similarities and differences with coral reefs, but can all be placed within the same geometry defined by the dimensions of competition and growth. The spatial Lotka-Volterra model that the geometry represents is also testable using data from these diverse ecosystems, because it makes specific predictions about when and what kind of shifts should occur as competition and growth ratios vary. Such a cross-system empirical synthesis can potentially facilitate the exchange of diverse conservation experiences. Moreover, the geometry highlights that regime shifts (Scheffer and Carpenter 2003) should be considered more broadly to include transitions between coexistence and single-species dominance, rather than being solely associated with alternative stable states. Coral reefs (Hughes et al. 2017, Darling et al. 2019) and other ecosystems (Waters et al. 2016) face multiple stressors and perturbations simultaneously in the Anthropocene, resulting in challenging complexities unless ecological theory sheds light on their commonalities and interactions. The geometric perspective is one potential tool to distill complexity, avoid simplistic explanations, and facilitate multiple management options for conservation success.

## Acknowledgements

We thank Tim Essington, Kevin McCann, Peter Mumby, Steve Palumbi, and Daniel Schindler for discussions. Research was supported by the Gordon and Betty Moore Foundation, the Nature Conservancy, a Canadian Natural Sciences and Engineering Research Council Scholarship (CGS-D), and US National Science Foundation awards OCE-1426891 and DEB-1616821.

## Author’s Contributions

EWT, LCM, AG, MAC, MSW, and MLP conceptualized the project, EWT and AG conducted the analyses, and all authors contributed to writing.

## Appendix

### Lotka-Volterra Model

The stability of an equilibrium set is indicated by whether the eigenvalues of the Jacobian matrix are negative. The eigenvalues for the first case of single-species equilibrium (Table A1), with only species *i* surviving, are:

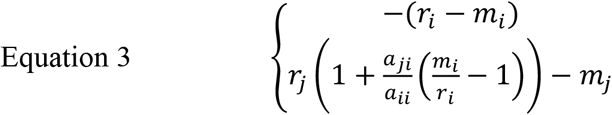

The first line is always negative, so the sufficient and necessary condition for species *i* stability hinges on line two, which translates to the inequality in Equation 1.

### Coral-Macroalgal Model

Mumby et al.’s model (Mumby et al. 2007) of coral cover (*N_1_*), algal turf cover (*T*), and macroalgae (*N_2_*) contains five parameters: coral birth rate (*r*), coral mortality (*d*), macroalgal birth rate (*γ*), macroalgal overgrowth rate on coral (*a*), and grazing rate (*g*). The model consists of three equations:

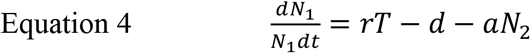

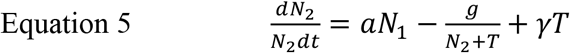

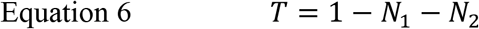

Turf is simply empty space from the perspective of corals and macroalgae. The solutions are:

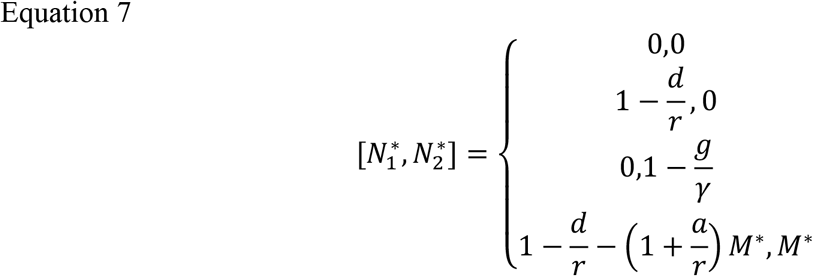

where

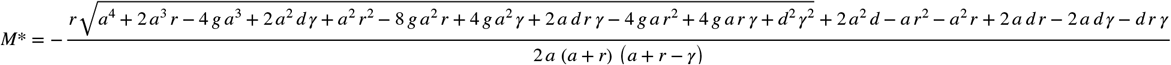

The dynamic equations for *N_1_* and *N_2_* can be written in Lotka-Volterra form. First, the growth of coral is:

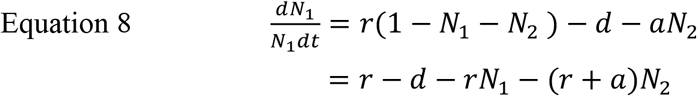

Clearly, interspecific competition with macroalgae (-*r-a*) is stronger than intraspecific competition within coral (-*r*). In this form, it is clear that the interactions modelled are predation, competition for empty space, and grazing. The term *a* is an antisymmetric predator-prey (+/-) interaction effect between macroalgae and corals.

Second, the growth of macroalgae is:

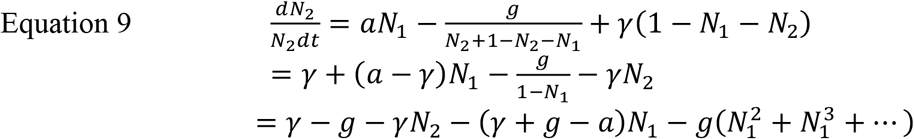

The interaction is negative for macroalgae through grazing (*g*) but can be positive when *a*>*γ*+*g* and *N_1_* is low. The negative effect of coral on macroalgae is amplified at increasing *N_1_* (through the Taylor series). In the simplified case of *a*=*γ*, when macroalgae overgrows corals and turf at the same rate, the interaction with coral is simply -*g*/(1-*N_1_*) or -*g*(1+*N*_1_+*N_1_^2^*+*N_1_^3^*+…) according to the geometric power series (when |*N_1_*|<1), which is increasingly negative as *N_1_* increases. In general, interspecific competition can be stronger than intraspecific competition when *g>a*, even without higher order terms; this becomes even more likely with higher order terms.

For macroalgae, *γ* is a spatial competition rate among themselves and with corals, whereas for corals, *r* is the analogous spatial competition rate. Additionally, macroalgae is removed by corals at a rate proportional to 1/(1-*N_1_*), although corals do not directly benefit from this process.

If we drop the nonlinear terms (*N_1_^2^*+*N_1_^3^*+…), the equilibria are, according to the Lotka-Volterra solutions (Table 1):

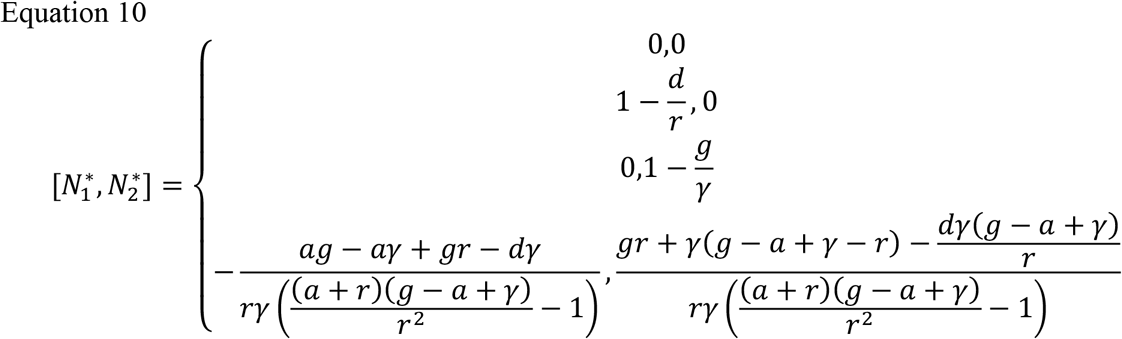

### Spatial Model

The term *C_ij_N_j_* is the local density of species *j* around species *i*, and can also be written as *N_j_*+*c_ij_/N_i_* where *C_ij_* is the average spatial covariance weighted by an interaction kernel (Bolker and Pacala 1999). Thus, *C_ij_* = *1+c_ij_*/(*N_i_N_j_*). Note by definition *C_12_=C_21_*. Assuming interactions only occur within a fixed local area (an interaction kernel that takes the value of 1 within the local area, and 0 everywhere else), *c_ij_=*E[(*n_i_-N_i_*)(*n_j_-N_j_*)], where *n_i_* is the number of individuals of species *i* at a location. For *i=j*, *C_ij_* is just the spatial variance in the number of *i* individuals.

Consider the simplified symmetric case where *m_i_*=0, *C_11_=C_22_*, *a_11_*=*a_22_*, and *a_12_*=*a_21_*. Then, the total density at coexistence is:

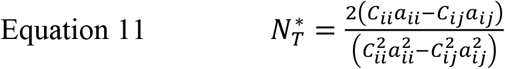

An increase in intraspecific clustering (*C_ij_*) and a decrease in interspecific clustering (*C_ij_*), as expected with decreased dispersal or connectivity (Bolker and Pacala 1999), would cause *N_T_^*^* to decrease. However, this total density can still be greater than the non-clustering single-species population if:

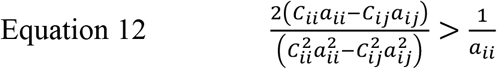

This condition simplifies to:

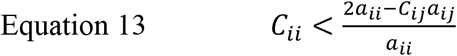

Thus, intraspecific clustering should be relatively small for the stable coexisting community to be denser than a single-species population (a positive diversity effect). A sufficient but not necessary condition is *C_ii_*<2 (obtained by assuming complete segregation between species, *C_ij_*=0). On the other hand, the stable coexistence condition in this simplified symmetric example is (reverse of Equation 2 where the right-hand-side equals 1):

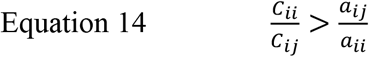

This condition states that intraspecific clustering should be large relative to interspecific clustering for stable coexistence. We obtain the condition (Equation 15) for a community’s total density to be greater than a non-clustering single-species population by joining Equation 13 and Equation 14.

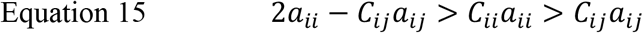

The relationship between interspecific and intraspecific clustering (*C_ij_* vs. *C_ii_*) can be complicated. The ratio *C_ij_*/*C_ii_* can be derived exactly for two-player games on graphs assuming constant total population size (Matsuda et al. 1992, Nathanson et al. 2009), but only approximately for population dynamics in continuous space (Bolker and Pacala 1997, 1999) as a function of growth and movement rates. The latter framework and other simulations (Tekwa et al. 2019b) show that intraspecific clustering characteristically increases with decreased movement rate, while interspecific clustering decreases at a comparatively slower rate with decreased movement rate. Thus, we assumed a characteristic relationship *C_ij_*=*C_ii_^-3^*, which creates the three relative clustering levels [1, 2, 4] and corresponding intra-[1, 1.19, 1.41] and interspecific clustering [1, 0.59, 0.35] used for the spatial clustering bifurcation (Table 2, Figure 2, and Figure 3).

### Appendix Figures

**Figure A1.**
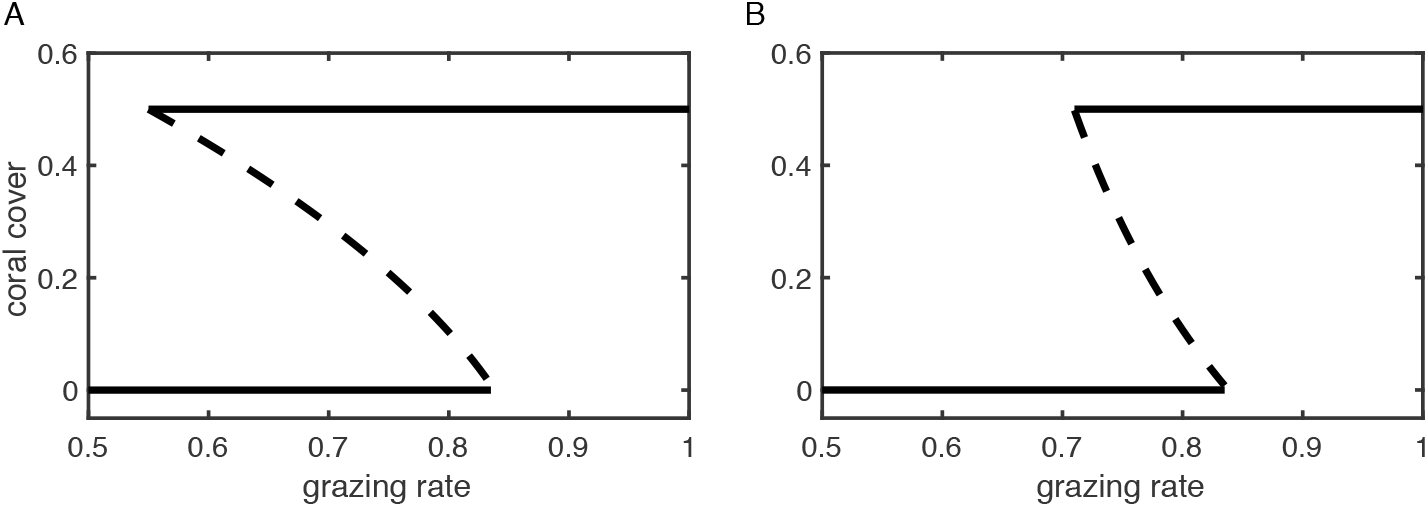
Coral-macroalgal model solutions. **(A)** The original Mumby model with non-linear coral competition affecting macroalgae. **(B)** A linearized Lotka-Volterra version of the Mumby model. The solid lines are analytical stable states, and the dashed curves are the unstable saddle-nodes.

### Appendix Tables

**Table A1.** Model equilibria. All symbols are defined in Table 2, and *M*\* (coexistence macroalgal cover) is given in Equation 7.

| Model | Species (i) | Equilibrium 1 ($N_i^*$) | Equilibrium 2 ($N_i^*$) | Coexistence equilibrium ($N_i^*$) |
| --- | --- | --- | --- | --- |
| Lotka-Volterra | 1 | $\frac{r_1 - m_1}{r_1 a_{11}}$ | 0 | $\frac{r_1 r_2 (a_{22} - a_{12}) + a_{12} m_2 r_1 - a_{22} m_1 r_2}{r_1 r_2 (a_{11} a_{22} - a_{12} a_{21})}$ |
| | 2 | 0 | $\frac{r_2 - m_2}{r_2 a_{22}}$ | $\frac{r_1 r_2 (a_{11} - a_{21}) + a_{21} m_1 r_2 - a_{11} m_2 r_1}{r_1 r_2 (a_{11} a_{22} - a_{12} a_{21})}$ |
| Mumby | 1 Coral | $1 - \frac{d}{r}$ | 0 | $1 - \frac{d}{r} - \left(1 + \frac{a}{r}\right) M^*$ |
| | 2 Algae | 0 | $1 - \frac{g}{\gamma}$ | $M^*$ |
| Spatial Lotka-Volterra | 1 | $\frac{r_1 - m_1}{r_1 a_{11} C_{11}}$ | 0 | $\frac{r_1 r_2 (a_{22} C_{22} - a_{12} C_{12}) + a_{12} C_{12} m_2 r_1 - a_{22} C_{22} m_1 r_2}{r_1 r_2 (a_{11} C_{11} a_{22} C_{22} - a_{12} a_{21} C_{12}^2)}$ |
| | 2 | 0 | $\frac{r_2 - m_2}{r_2 a_{22} C_{22}}$ | $\frac{r_1 r_2 (a_{11} C_{11} - a_{21} C_{12}) + a_{21} C_{12} m_1 r_2 - a_{11} C_{11} m_2 r_1}{r_1 r_2 (a_{11} C_{11} a_{22} C_{22} - a_{12} a_{21} C_{12}^2)}$ |

**Table A2.**
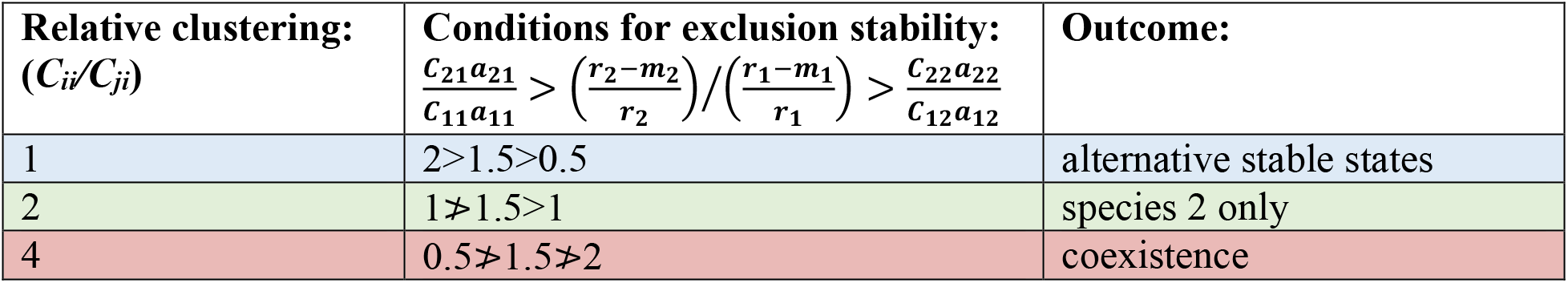
Effect of clustering on two-species community outcomes. Parameters are: *a_ii_/a_ji_*=0.5 (intra-to-interspecific cross-competition ratio) and ((*r_2_-m_2_*)/*r_2_*)/((*r_1_-m_1_*)/*r_1_*)=1.5 (intrinsic growth inequality). Relative clustering is defined as intra-to-interspecific clustering ratio (*C_ii_/C_ji_*).

**Table A3.**
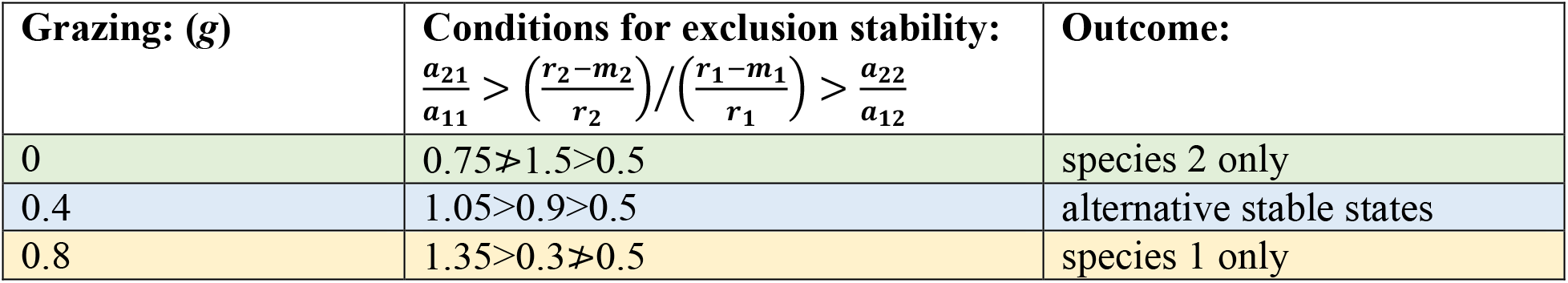
Effect of grazing on two-species community outcomes. Parameters are: *a_11_/a_21_*=1.33/(1+*g*), *a_22_/a_12_*=0.5 (intra-to-interspecific cross-competition ratios), and ((*r_2_-m_2_*)/*r_2_*)/((*r_1_-m_1_*)/*r_1_*)=(1-*g*)/0.66 (intrinsic growth inequality). Species 1 and 2 correspond to coral and macroalgae, respectively. Grazing rate on macroalgae is *g*.

**Table A4.** Numerical outcomes of grazing bifurcations. Low and high grazing rates under the scenarios of no warming, warming, and a range of relative clustering (corresponding to Figure 2 and Figure 3), as well as their corresponding parameters in competition (*α_1_*,*α_2_*) and intrinsic growth (*f_21_*) terms. Regime outcomes for parameter sets are shown in color (yellow=species 1 or coral only, green=species 2 or macroalgae only, red=coexistence).

| **Scenario** | low grazing ($g=0.55$) | | | high grazing ($g=0.85$) | | |
| --- | --- | --- | --- | --- | --- | --- |
| +0°C | $\log_2(\alpha_1)$ | $\log_2(\alpha_2)$ | $\log_2(f_{21})$ | $\log_2(\alpha_1)$ | $\log_2(\alpha_2)$ | $\log_2(f_{21})$ |
| **(A)** $C_{ii}/C_{ij}=1$ | 1 | -1.1 | 0 | 0.37 | -1.1 | -1.1 |
| **(B)** $C_{ii}/C_{ij}=2$ | 2 | -0.070 | 0 | 1.4 | -0.070 | -1.1 |
| **(C)** $C_{ii}/C_{ij}=4$ | 3 | 0.93 | 0 | 2.4 | 0.93 | -1.1 |
| +1°C | $\log_2(\alpha_1)$ | $\log_2(\alpha_2)$ | $\log_2(f_{21})$ | $\log_2(\alpha_1)$ | $\log_2(\alpha_2)$ | $\log_2(f_{21})$ |
| **(D)** $C_{ii}/C_{ij}=1$ | 0.64 | -1.3 | 1.0 | 0.011 | -1.3 | -4.5 |
| **(E)** $C_{ii}/C_{ij}=2$ | 1.6 | -0.27 | 1.0 | 1.0 | -0.27 | -4.5 |
| **(F)** $C_{ii}/C_{ij}=4$ | 2.6 | 0.73 | 1.0 | 2.0 | 0.73 | -4.5 |

## References

Anthony, K. R. N., J. A. Maynard, G. Diaz-Pulido, P. J. Mumby, P. A. Marshall, L. Cao, and O. Hoegh-Guldberg. 2011. Ocean acidification and warming will lower coral reef resilience: CO2 and coral reef resilience. Global Change Biology 17:1798–1808.

Baskett, M. L., N. S. Fabina, and K. Gross. 2014. Response Diversity Can Increase Ecological Resilience to Disturbance in Coral Reefs. The American Naturalist 184:E16–E31.

Blackwood, J. C., A. Hastings, and P. J. Mumby. 2011. A model-based approach to determine the long-term effects of multiple interacting stressors on coral reefs. Ecological Applications 21:2722–2733.

Bolker, B. M., and S. W. Pacala. 1999. Spatial Moment Equations for Plant Competition: Understanding Spatial Strategies and the Advantages of Short Dispersal. American Naturalist 153:575–602.

Bolker, B., and S. W. Pacala. 1997. Using Moment Equations to Understand Stochastically Driven Spatial Pattern Formation in Ecological Systems. Theoretical Population Biology 52:179–197.

Bomze, I. M. 1983. Lotka-Volterra equation and replicator dynamics: A two-dimensional classification. Biological Cybernetics 48:201–211.

Botsford, L. W., J. C. Castilla, and C. H. Peterson. 1997. The Management of Fisheries and Marine Ecosystems. Science 277:509–515.

Bruno, J. F., H. Sweatman, W. F. Precht, E. R. Selig, and V. G. W. Schutte. 2009. Assessing evidence of phase shifts from coral to macroalgal dominance on coral reefs. Ecology 90:1478–1484.

Chesson, P. 2000. General Theory of Competitive Coexistence in Spatially-Varying Environments. Theoretical Population Biology 58:211–237.

Costello, C., D. Ovando, T. Clavelle, C. K. Strauss, R. Hilborn, M. C. Melnychuk, T. A. Branch, S. D. Gaines, C. S. Szuwalski, R. B. Cabral, D. N. Rader, and A. Leland. 2016. Global fishery prospects under contrasting management regimes. Proceedings of the National Academy of Sciences 113:5125–5129.

Darling, E. S., T. R. McClanahan, J. Maina, G. G. Gurney, N. A. J. Graham, F. Januchowski-Hartley, J. E. Cinner, C. Mora, C. C. Hicks, E. Maire, M. Puotinen, W. J. Skirving, M. Adjeroud, G. Ahmadia, R. Arthur, A. G. Bauman, M. Beger, M. L. Berumen, L. Bigot, J. Bouwmeester, A. Brenier, T. C. L. Bridge, E. Brown, S. J. Campbell, S. Cannon, B. Cauvin, C. A. Chen, J. Claudet, V. Denis, S. Donner, Estradivari, N. Fadli, D. A. Feary, D. Fenner, H. Fox, E. C. Franklin, A. Friedlander, J. Gilmour, C. Goiran, J. Guest, J.-P. A. Hobbs, A. S. Hoey, P. Houk, S. Johnson, S. D. Jupiter, M. Kayal, C. Kuo, J. Lamb, M. A. C. Lee, J. Low, N. Muthiga, E. Muttaqin, Y. Nand, K. L. Nash, O. Nedlic, J. M. Pandolfi, S. Pardede, V. Patankar, L. Penin, L. Ribas-Deulofeu, Z. Richards, T. E. Roberts, K. S. Rodgers, C. D. M. Safuan, E. Sala, G. Shedrawi, T. M. Sin, P. SmallhornWest, J. E. Smith, B. Sommer, P. D. Steinberg, M. Sutthacheep, C. H. J. Tan, G. J. Williams, S. Wilson, T. Yeemin, J. F. Bruno, M.-J. Fortin, M. Krkosek, and D. Mouillot. 2019. Social–environmental drivers inform strategic management of coral reefs in the Anthropocene. Nature Ecology & Evolution.

Dudgeon, S., R. Aronson, J. Bruno, and W. Precht. 2010. Phase shifts and stable states on coral reefs. Marine Ecology Progress Series 413:201–216.

Durrett, R., and S. Levin. 1994. The Importance of Being Discrete (and Spatial). Theoretical Population Biology 46:363–394.

Elmhirst, T., S. R. Connolly, and T. P. Hughes. 2009. Connectivity, regime shifts and the resilience of coral reefs. Coral Reefs 28:949–957.

Fabina, N. S., M. L. Baskett, and K. Gross. 2015. The differential effects of increasing frequency and magnitude of extreme events on coral populations. Ecological Applications 25:1534–1545.

Gibbs, J. W. 1873. A Method of Geometrical Representation of the Thermodynamic Properties by Means of Surfaces. Transactions of Connecticut Academy of Arts and Sciences:382–404.

Gordon, H. S. 1954. The Economic Theory of a Common-Property Resource: The Fishery. The Journal of Political Economy 62:124–142.

Graham, N. A. J., S. Jennings, M. A. MacNeil, D. Mouillot, and S. K. Wilson. 2015. Predicting climate-driven regime shifts versus rebound potential in coral reefs. Nature 518:94–97.

Hirota, M., M. Holmgren, E. H. Van Nes, and M. Scheffer. 2011. Global Resilience of Tropical Forest and Savanna to Critical Transitions. Science 334:232–235.

Hughes, T. P., M. L. Barnes, D. R. Bellwood, J. E. Cinner, G. S. Cumming, J. B. C. Jackson, J. Kleypas, I. A. van de Leemput, J. M. Lough, T. H. Morrison, S. R. Palumbi, E. H. van Nes, and M. Scheffer. 2017. Coral reefs in the Anthropocene. Nature 546:82–90.

Hughes, T. P., J. T. Kerry, S. R. Connolly, A. H. Baird, C. M. Eakin, S. F. Heron, A. S. Hoey, M. O. Hoogenboom, M. Jacobson, G. Liu, M. S. Pratchett, W. Skirving, and G. Torda. 2019. Ecological memory modifies the cumulative impact of recurrent climate extremes. Nature Climate Change 9:40–43.

Jiang, J., Z.-G. Huang, T. P. Seager, W. Lin, C. Grebogi, A. Hastings, and Y.-C. Lai. 2018. Predicting tipping points in mutualistic networks through dimension reduction. Proceedings of the National Academy of Sciences 115:E639–E647.

Jokiel, P. L., K. S. Rodgers, E. K. Brown, J. C. Kenyon, G. Aeby, W. R. Smith, and F. Farrell. 2015. Comparison of methods used to estimate coral cover in the Hawaiian Islands. PeerJ 3:e954.

Knowlton, N. 1992. Thresholds and Multiple Stable States in Coral Reef Community Dynamics. American Zoologist 32:674–682.

van de Leemput, I. A., T. P. Hughes, E. H. van Nes, and M. Scheffer. 2016. Multiple feedbacks and the prevalence of alternate stable states on coral reefs. Coral Reefs 35:857–865.

Li, X., H. Wang, Z. Zhang, and A. Hastings. 2014. Mathematical analysis of coral reef models. Journal of Mathematical Analysis and Applications 416:352–373.

Ling, S. D., R. E. Scheibling, A. Rassweiler, C. R. Johnson, N. Shears, S. D. Connell, A. K. Salomon, K. M. Norderhaug, A. Perez-Matus, J. C. Hernandez, S. Clemente, L. K. Blamey, B. Hereu, E. Ballesteros, E. Sala, J. Garrabou, E. Cebrian, M. Zabala, D. Fujita, and L. E. Johnson. 2014. Global regime shift dynamics of catastrophic sea urchin overgrazing. Philosophical Transactions of the Royal Society B: Biological Sciences 370:20130269–20130269.

Lion, S., and M. van Baalen. 2008. Self-structuring in spatial evolutionary ecology. Ecology Letters 11:277–295.

Lotka, A. J. 1978. The growth of mixed populations: Two species competing for a common food supply. Pages 274–286 The Golden Age of Theoretical Ecology: 1923–1940. Springer Berlin Heidelberg, Berlin, Heidelberg.

Matsuda, H., N. Ogita, A. Sasaki, and K. Sato. 1992. Statistical Mechanics of Population: The Lattice Lotka-Volterra Model. Progress of Theoretical Physics 88:1035–1049.

Maxwell, J. C., and P. M. Harman. 1990. The scientific letters and papers of James Clerk Maxwell. Cambridge University Press, Cambridge [England]; New York.

McCann, K., and P. Yodzis. 1995. Bifurcation Structure of a Three-Species Food-Chain Model. Theoretical Population Biology 48:93–125.

McCook, L., J. Jompa, and G. Diaz-Pulido. 2001. Competition between corals and algae on coral reefs: a review of evidence and mechanisms. Coral Reefs 19:400–417.

McManus, L. C., J. R. Watson, V. V. Vasconcelos, and S. A. Levin. 2019. Stability and recovery of coral-algae systems: the importance of recruitment seasonality and grazing influence. Theoretical Ecology 12:61–72.

Mumby, P. J., A. Hastings, and H. J. Edwards. 2007. Thresholds and the resilience of Caribbean coral reefs. Nature 450:98–101.

Mumby, P. J., R. S. Steneck, and A. Hastings. 2013. Evidence for and against the existence of alternate attractors on coral reefs. Oikos 122:481–491.

Nathanson, C. G., C. E. Tarnita, and M. A. Nowak. 2009. Calculating Evolutionary Dynamics in Structured Populations. PLoS Computational Biology 5:e1000615.

Neuhauser, C., and S. W. Pacala. 1999. An explicitly spatial version of the Lotka-Volterra model with interspecific competition. The Annals of Applied Probability 9:1226–1259.

Petraitis, P., and C. Hoffman. 2010. Multiple stable states and relationship between thresholds in processes and states. Marine Ecology Progress Series 413:189–200.

Sandin, S. A., and D. E. McNamara. 2012. Spatial dynamics of benthic competition on coral reefs. Oecologia 168:1079–1090.

Scheffer, M., S. Carpenter, J. A. Foley, C. Folke, and B. Walker. 2001. Catastrophic shifts in ecosystems. Nature 413:591–596.

Scheffer, M., and S. R. Carpenter. 2003. Catastrophic regime shifts in ecosystems: linking theory to observation. Trends in Ecology & Evolution 18:648–656.

Scheffer, M., S. H. Hosper, M.-L. Meijer, B. Moss, and E. Jeppesen. 1993. Alternative equilibria in shallow lakes. Trends in Ecology & Evolution 8:275–279.

Schmitt, R. J., S. J. Holbrook, S. L. Davis, A. J. Brooks, and T. C. Adam. 2019. Experimental support for alternative attractors on coral reefs. Proceedings of the National Academy of Sciences 116:4372–4381.

Schröder, A., L. Persson, and A. M. De Roos. 2005. Direct experimental evidence for alternative stable states: a review. Oikos 110:3–19.

Solow, R. M. 1956. A Contribution to the Theory of Economic Growth. The Quarterly Journal of Economics 70:65.

Staver, A. C., and S. A. Levin. 2012. Integrating Theoretical Climate and Fire Effects on Savanna and Forest Systems. The American Naturalist 180:211–224.

Steffen, W., K. Richardson, J. Rockstrom, S. E. Cornell, I. Fetzer, E. M. Bennett, R. Biggs, S. R. Carpenter, W. de Vries, C. A. de Wit, C. Folke, D. Gerten, J. Heinke, G. M. Mace, L. M. Persson, V. Ramanathan, B. Reyers, and S. Sorlin. 2015. Planetary boundaries: Guiding human development on a changing planet. Science 347:1259855–1259855.

Tarnita, C. E., H. Ohtsuki, T. Antal, F. Fu, and M. A. Nowak. 2009. Strategy selection in structured populations. Journal of Theoretical Biology 259:570–581.

Tekwa, E. W., E. P. Fenichel, S. A. Levin, and M. L. Pinsky. 2019a. Path-dependent institutions drive alternative stable states in conservation. Proceedings of the National Academy of Sciences 116:689–694.

Tekwa, E. W., A. Gonzalez, and M. Loreau. 2015. Local densities connect spatial ecology to game, multilevel selection and inclusive fitness theories of cooperation. Journal of Theoretical Biology 380:414–425.

Tekwa, E. W., A. Gonzalez, and M. Loreau. 2019b. Spatial evolutionary dynamics produce a negative cooperation–population size relationship. Theoretical Population Biology 125:94–101.

Tekwa, E. W., D. Nguyen, M. Loreau, and A. Gonzalez. 2017. Defector clustering is linked to cooperation in a pathogenic bacterium. Proceedings of the Royal Society B: Biological Sciences 284:20172001.

Tilman, D. 1980. Resources: A Graphical-Mechanistic Approach to Competition and Predation. The American Naturalist 116:362–393.

Verwiebe, F. L. 1939. A P-V-T Diagram of the Allotropic Forms of Ice. American Journal of Physics 7:187–189.

Volterra, V. 1926. Variations and fluctuations of the number of individuals in animal species living together. Animal ecology:409–448.

van der Waals, J. D. 1873. Over de Continuiteit van den Gasen Vloeistoftoestand. Leiden.

Waters, C. N., J. Zalasiewicz, C. Summerhayes, A. D. Barnosky, C. Poirier, A. Ga uszka, A. Cearreta, M. Edgeworth, E. C. Ellis, M. Ellis, C. Jeandel, R. Leinfelder, J. R. McNeill, D. d. Richter, W. Steffen, J. Syvitski, D. Vidas, M. Wagreich, M. Williams, A. Zhisheng, J. Grinevald, E. Odada, N. Oreskes, and A. P. Wolfe. 2016. The Anthropocene is functionally and stratigraphically distinct from the Holocene. Science 351:aad2622.

